# General sexual desire, but not desire for uncommitted sexual relationships, tracks changes in women’s hormonal status

**DOI:** 10.1101/155788

**Authors:** Benedict C Jones, Amanda C Hahn, Claire I Fisher, Hongyi Wang, Michal Kandrik, Lisa M DeBruine

**Affiliations:** Institute of Neuroscience & Psychology, University of Glasgow, UK.; Department of Psychology, Humboldt State University, USA.

**Keywords:** sexual desire, sociosexuality, progesterone, estradiol, testosterone, mating psychology

## Abstract

Several recent longitudinal studies have investigated the hormonal correlates of both young adult women’s general sexual desire and, more specifically, their desire for uncommitted sexual relationships. Findings across these studies have been mixed, potentially because each study tested only small samples of women (Ns = 43, 33, and 14). Here we report results from a much larger (N = 375) longitudinal study of hormonal correlates of young adult women’s general sexual desire and their desire for uncommitted sexual relationships. Our analyses suggest that within-woman changes in general sexual desire are negatively related to progesterone, but are not related to testosterone or cortisol. We observed some positive relationships for estradiol, but these were generally only significant for solitary sexual desire. By contrast with our results for general sexual desire, analyses showed no evidence that changes in women’s desire for uncommitted sexual relationships are related to their hormonal status. Together, these results suggest that changes in hormonal status contribute to changes in women’s general sexual desire, but do not influence women’s desire for uncommitted sexual relationships.

## 1. Introduction

Links between within-subject changes in steroid hormone levels and sexual desire in circum-menopausal and post-menopausal women have been extensively studied (reviewed in Cappelletti & Wallen, 2016 and Motta-Mena & Puts, 2017). While it is well established that sexual desire varies across the menstrual cycle in young adult women (reviewed in Motta-Mena & Puts, 2017 and Roney & Simmons, 2013), surprisingly little is known about the specific hormonal correlates of within-subject changes in young adult women’s sexual desire (Grebe et al., 2016; Motta-Mena & Puts, 2017; Roney & Simmons, 2013, 2016; Wallen, 2013).

To directly address this issue, Roney and Simmons (2013) used a longitudinal design to investigate possible relationships between salivary estradiol, progesterone, and testosterone and self-ratings of general sexual desire in a sample of 43 women. Their analyses suggested a positive effect of estradiol, a negative effect of progesterone, and no effect of testosterone on general sexual desire.

Grebe et al. (2016) reported similar analyses for a sample of 33 women in romantic relationships. By contrast with Roney and Simmons’ (2013) results, Grebe et al. (2016) reported a negative effect of estradiol and no effects of progesterone or testosterone on general sexual desire. Note that the effects of estradiol in Grebe et al. (2016) and Roney and Simmons (2013) were in opposite directions.

Grebe et al. (2016) suggested that these strikingly different results could occur if hormones have different effects on women’s general sexual desire and their desire for uncommitted sexual relationships. Consistent with this explanation, they reported that estradiol had a positive effect and progesterone had a negative effect on the extent to which women in romantic relationships reported greater desire for extra-pair sex (i.e., sex with men other than their romantic partner) over in-pair sex (i.e., sex with their romantic partner). However, Roney and Simmons (2016) did not replicate Grebe et al’s (2016) results in a sample of 14 women in romantic relationships. Instead, they found that progesterone had negative effects on both extra-pair and in-pair sexual desire, suggesting that progesterone simply has a negative effect on general sexual desire.

In summary, despite several recent studies on the topic, the relationships between changes in women’s hormonal status and changes in their general sexual desire and desire for uncommitted sexual relationships remain unclear. One potentially important limitation of previous studies is that they tested only small samples of women (N=33, each woman tested twice, Grebe et al., 2016; N=43, each woman tested >14 times, Roney & Simmons, 2013; N=14, each woman tested >14 times, Roney & Simmons, 2016). In light of this issue, here we report results from a much larger longitudinal study of the hormonal correlates of women’s general sexual desire and their desire for uncommitted sexual relationships (N=375).

## 2. Methods

### 2.1. Participants

We tested 375 heterosexual women (mean age=21.56 years, SD=3.31 years) who reported that they were not using any form of hormonal contraceptive (i.e., reported having natural menstrual cycles). Participants completed up to three blocks of test sessions. Each of the three blocks of test sessions consisted of five weekly test sessions. Women participated as part of a large study of possible effects of steroid hormones on women’s behavior (Jones et al., 2017a). The data analyzed here are all responses from blocks of test sessions where women were not using any form of hormonal contraceptive and provided data for at least one of the measures of sexual desire or sociosexual orientation. So that results could be directly compared with the data Roney and Simmons (2013, 2016) and Grebe et al. (2016) reported, only responses from blocks of test sessions where women were not using any form of hormonal contraceptive were analyzed in the current study. Following these restrictions, 337 women had completed five or more test sessions and 98 of these women completed ten test sessions. Thirty-eight women completed fewer than five test sessions.

### 2.2. Procedure

In each test session, women reported their current romantic partnership status (partnered or unpartnered), provided a saliva sample, and completed Spector et al’s (1996) Sexual Desire Inventory (SDI-2), a rating of current sex drive, and Penke and Asendorpf’s (2008) Revised Sociosexual Orientation Inventory (SOI-R). The SDI-2 and rating of current sex drive assess general sexual desire, while subscales of the SOI-R assess desire for (and attitudes to) uncommitted sexual relationships. Questionnaire order was fully randomized.

The Sexual Desire Inventory (SDI-2) is a 14-item questionnaire that assesses general sexual desire (Spector et al., 1996). An example question is “When you are in romantic situations (such as a candle lit dinner, a walk on the beach, etc.), how strong is your sexual desire?”, to which participants responded using a 1 (no desire) to 9 (strong desire) scale. As well as providing a score for total sexual desire (M=44.15, SD=15.66), the SDI-2 also provides separate scores for desire for sexual activity with another person (dyadic sexual desire, M=35.51, SD=11.95) and desire for sexual activity by oneself (solitary sexual desire, M=8.63, SD=6.46).

Women also rated their current sex drive on a 1 (very low) to 7 (very high) scale. This question is similar to the single item used to assess general sexual desire in Roney and Simmons (2013). Each woman answered this question twice in each test session. Their reported current sex drive score for each test session was the average of these two ratings (M=3.77, SD=1.56).

The Revised Sociosexual Orientation Inventory (SOI-R) is a nine-item questionnaire that assesses openness to uncommitted sexual relationships (Penke & Asendorpf, 2008). Each item is answered using a 1 to 5 scale. The SOI-R has three components (desire, attitude, and behavior). The desire component consists of 3 items (e.g., “In everyday life, how often do you have spontaneous fantasies about having sex with someone you have just met?”), for which 1 on the response scale corresponds to “never” and 5 corresponds to “nearly every day” (M=8.06, SD=2.96). The attitude component consists of 3 items (e.g., “Sex without love is OK”), for which 1 on the response scale corresponds to “totally disagree” and 5 corresponds to “totally agree” (M=9.22, SD=3.50). The behavior component consists of 3 items (e.g., “With how many different partners have you had sex within the past 12 months?”), for which 1 on the response scale corresponds to “0 sexual partners” and 5 corresponds to “8 or more sexual partners” (M=5.74, SD=2.67). Scores for each component are calculated by summing the individual scores for the 3 relevant items.

### 2.3. Saliva samples

Participants provided a saliva sample via passive drool (Papacosta & Nassis, 2011) in each test session. Participants were instructed to avoid consuming alcohol and coffee in the 12 hours prior to participation and avoid eating, smoking, drinking, chewing gum, or brushing their teeth in the 60 minutes prior to participation. Each woman’s test sessions took place at approximately the same time of day to minimize effects of diurnal changes in hormone levels (Veldhuis et al., 1988; Bao et al., 2003).

Saliva samples were frozen immediately and stored at −32°C until being shipped, on dry ice, to the Salimetrics Lab (Suffolk, UK) for analysis, where they were assayed using the Salivary 17β-Estradiol Enzyme Immunoassay Kit 1-3702 (M=3.30 pg/mL, SD=1.27 pg/mL, sensitivity=0.1 pg/mL, intra-assay CV=7.13%, inter-assay CV=7.45%), Salivary Progesterone Enzyme Immunoassay Kit 1-1502 (M=148.59 pg/mL, SD=96.22 pg/mL, sensitivity=5 pg/mL, intra-assay CV=6.20%, inter-assay CV=7.55%), Salivary Testosterone Enzyme Immunoassay Kit 1-2402 (M=87.57 pg/mL, SD=27.19 pg/mL, sensitivity<1.0 pg/mL, intra-assay CV=4.60%, inter-assay CV=9.83%), and Salivary Cortisol Enzyme Immunoassay Kit 1-3002 (M=0.23 μg/dL, SD=0.16 μg/dL, sensitivity<0.003 μg/dL, intra-assay CV=3.50%, inter-assay CV=5.08%). Although Roney and Simmons (2013, 2016) and Grebe et al. (2016) did not consider possible effects of cortisol in their studies, we included cortisol in our study because some studies suggest links between cortisol and women’s attractiveness judgments of potential mates (e.g. Ditzen et al., 2017).

Hormone levels more than three standard deviations from the sample mean for that hormone or where Salimetrics indicated levels were outside the sensitivity range of their relevant ELISA were excluded from the dataset (~1 % of hormone measures were excluded for these reasons). The descriptive statistics given above do not include these excluded values. Values for each hormone were centered on their subject-specific means to isolate effects of within-subject changes in hormones. They were then scaled so the majority of the distribution for each hormone varied from −.5 to .5 to facilitate calculations in the linear mixed models. Since hormone levels were centered on their subject-specific means, women with only one value for a hormone could not be included in these analyses.

### 2.4. Analyses

Linear mixed models were used to test for possible effects of hormonal status on sexual desire and sociosexuality. Analyses were conducted using R version 3.3.2 (R Core Team, 2016), with lme4 version 1.1-13 (Bates et al., 2014) and lmerTest version 2.0-33 (Kuznetsova et al., 2013). The dependent variable was questionnaire or subscale score (separate models were run for each questionnaire or subscale). Predictors were scaled and centered hormone levels. Random slopes were specified maximally following Barr et al. (2013) and Barr (2013). Full model specifications and full results for each analysis are given in our Supplemental Information. Data files and analysis scripts are publicly available at https://osf.io/8bph4/.

## 3. Results

Scores for each questionnaire or subscale were analyzed separately. For each dependent variable (i.e., questionnaire or subscale score) we ran three models. The first model (Model 1) included estradiol, progesterone, and their interaction as predictors. The second model (Model 2) included estradiol, progesterone, and estradiol-to-progesterone ratio as predictors. We tested for combined effects of estradiol and progesterone by including the estradiol by progesterone interaction (Model 1) and estradiol-to-progesterone ratio (Model 2) because both approaches have recently been used to test for combined effects of estradiol and progesterone in the hormones and behavior literature (see Puts et al., 2013 and Roney & Simmons, 2013 for examples of studies using one of these two approaches). The third model (Model 3) included only testosterone and cortisol as predictors. Our analysis strategy is identical to that used in Jones et al. (2017a) and Jones et al. (2017b) to investigate the hormonal correlates of women’s mate preferences and disgust sensitivity, respectively.

For each dependent variable, we also repeated the three models described above, this time including tests for possible moderating effects of women’s romantic partnership status (partnered versus unpartnered). None of the significant effects described below were qualified by higher-order interactions with partnership status, suggesting that they were not moderated by romantic partnership status. These additional analyses are reported in full in our Supplemental Information.

### 3.1. Total Sexual Desire (total score on SDI-2)

Model 1 revealed a significant negative effect of progesterone (estimate=−1.78, t=−2.56, p=.011). Neither the effect of estradiol (estimate=1.32, t=1.60, p=.11) nor the interaction between estradiol and progesterone (estimate=2.28, t=0.50, p=.62) were significant. Model 2 revealed a significant negative effect of progesterone (estimate=−2.43, t=−2.78, p=.006) and a significant positive effect of estradiol (estimate=1.73, t=2.05, p=.041). The effect of estradiol-to-progesterone ratio was not significant (estimate=−0.86, t=−1.60, p=.15). Model 3 showed no significant effects of either testosterone (estimate=−0.60, t=−0.72, p=.47) or cortisol (estimate=0.81, t=1.16, p=.25).

### 3.2. Dyadic Sexual Desire (score on dyadic subscale of SDI−2)

Model 1 revealed a significant negative effect of progesterone (estimate=−1.44, t=−2.48, p=.013). Neither the effect of estradiol (estimate=0.46, t=0.67, p=.50) nor the interaction between estradiol and progesterone (estimate=1.10, t=0.30, p=.76) were significant. Model 2 revealed a significant negative effect of progesterone (estimate=−1.97, t=−3.05, p=.002). A negative effect of estradiol-to-progesterone ratio was close to being significant (estimate=−0.64, t=−1.94, p=.052). The effect of estradiol was not significant (estimate=0.79, t=1.15, p=.25). Model 3 showed no significant effects of either testosterone (estimate=−0.88, t=−1.26, p=.21) or cortisol (estimate=0.63, t=1.09, p=.28).

### 3.3. Solitary Sexual Desire (score on solitary subscale of SDI−2)

Model 1 revealed a significant positive effect of estradiol (estimate=0.68, t=2.31, p=.021). Neither the effect of progesterone (estimate=−0.24, t=−0.96, p=.34) nor the interaction between estradiol and progesterone (estimate=0.63, t=0.41, p=.69) were significant. Model 2 revealed a significant positive effect of estradiol (estimate=0.69, t=2.31, p=.021). The effects of progesterone (estimate=−0.32, t=−0.98, p=.33) and estradiol-to-progesterone ratio (estimate=−0.17, t=−1. 19, p=.23) were not significant. Model 3 showed no significant effects of either testosterone (estimate=0.35, t=1.18, p=.24) or cortisol (estimate=0.03, t=0.14, p=.89).

### 3.4. Reported Current Sex Drive

Model 1 showed a negative effect of progesterone that was close to being significant (estimate=−0.27, t=−1.94, p=.052). Neither the effect of estradiol (estimate=0.06, t=0.33, p=.74) nor the interaction between estradiol and progesterone (estimate=−0.22, t=−0.23, p=.82) were significant. Model 2 revealed a significant negative effect of progesterone (estimate=−0.43, t=−2.35, p=.019). Neither the effect of estradiol (estimate=0.12, t=0.73, p=.47) nor the effect of estradiol-to-progesterone ratio (estimate=−0.13, t=−1.32, p=.20) was significant. Model 3 showed no significant effects of either testosterone (estimate=−0.07, t=−0.40, p=.69) or cortisol (estimate=0.21, t=1.62, p=. 11).

### 3.5. Sociosexual Desire (score on desire subscale of SOI-R)

Model 1 showed no significant effects of estradiol (estimate=0.28, t=1.55, p=.12), progesterone (estimate=−0.13, t=−0.88, p=.38), or their interaction (estimate=0.29, t=0.34, p=.74). Model 2 showed no significant effects of estradiol (estimate=0.26, t=1.38, p=. 17), progesterone (estimate=−0.05, t=−0.27, p=.79), or estradiol-to-progesterone ratio (estimate=0.08, t=0.83, p=.43). Model 3 showed no significant effects of either testosterone (estimate=0.02, t=0.08, p=.93) or cortisol (estimate=0.01, t=0.09, p=.93).

### 3.6. Sociosexual Attitude (score on attitude subscale of SOI-R)

Model 1 showed no significant effects of estradiol (estimate=0.01, t=0.06, p=.95), progesterone (estimate=−0.07, t=−0.47, p=.64), or their interaction (estimate=−0.46, t=−0.51, p=.61). Model 2 showed no significant effects of estradiol (estimate=0.07, t=0.34, p=.73), progesterone (estimate=−0.26, t=−1.44, p=.15), or estradiol-to-progesterone ratio (estimate=−0.19, t=−1.65, p=.12). Model 3 showed no significant effects of either testosterone (estimate=−0.20, t=−1.09, p=.28) or cortisol (estimate=−0.17, t=−1.27, p=.21).

### 3.7. Sociosexual Behavior (score on behavior subscale of SOI-R)

None of the questions on the behavior subscale of the SOI-R assess current sociosexuality (i.e., sociosexuality at the time of testing). Consequently, we did not analyze responses on the behavior subscale of the SOI-R. However, these behavior-subscale data, and the data for all our analyses, are publicly available at https://osf.io/8bph4/.

### 3.8. Additional analyses

Each of the significant effects described above remained significant when test-session order was included as a covariate. Note that this indicates that the significant hormonal effects that we observed could not be caused by questionnaire responses changing simply as a function of test-session order. They also remained significant when all nonsignificant independent variables were removed from the models. The effect of progesterone on total sexual desire was the only exception to this pattern. In that instance, the effect of progesterone was no longer significant (p=.092) when all other variables were removed from the models. Adding the interaction between testosterone and cortisol to models including those hormones did not reveal any significant effects of testosterone, cortisol, or their interaction for any of our dependent variables. These additional analyses are reported in full in our Supplemental Information.

## 4. Discussion

The current study was a large (N=375) longitudinal study of hormonal correlates of women’s general sexual desire and desire for uncommitted sexual relationships. Analyses of measures of women’s general sexual desire showed that progesterone had significant negative effects on total scores on the Sexual Desire Inventory (SDI−2), scores on the dyadic desire subscale of the SDI−2, and reported current sex drive. These results are consistent with Roney and Simmons (2016, 2013), who also reported a negative effect of progesterone on measures of women’s general sexual desire. Further analyses (see Supplemental Information) showed that none of these effects were moderated by women’s romantic partnership status.

Results of tests for effects of estradiol on measures of women’s general sexual desire were more mixed. Nonetheless, we observed positive effects of estradiol on the solitary desire subscale of the SDI−2 and, in one model (Model 2), total scores on the SDI−2. Again, further analyses (see Supplemental Information) showed that none of these effects were moderated by women’s romantic partnership status. These results provide some support for Roney and Simmons’ (2013) proposal that estradiol increases general sexual desire, although the effects of estradiol in our study were largely confined to the domain of solitary sexual desire.

Consistent with previous research (Grebe et al., 2016; Roney & Simmons, 2013, 2016), we found no evidence that measures of women’s general sexual desire were related to testosterone. Cortisol was also not related to women’s general sexual desire in our study, despite predicting women’s mate preferences in some recent work (Ditzen et al., 2017; but see Jones et al., 2017).

By contrast with our results for women’s general sexual desire, no hormones significantly predicted women’s responses on the desire or attitude subscales of the Revised Sociosexual Orientation Inventory (SOI-R). Although we did not assess women’s preferences for extra-pair versus in-pair sexual desire directly (Grebe et al., 2016), these results do not support the hypothesis that women’s desire for uncommitted sexual relationships is positively related to estradiol and negatively related to progesterone (Grebe et al., 2016). While the high test-retest reliability of the SOI-R subscales suggests they may measure trait-like aspects of sociosexuality (Penke & Asendorpf, 2008), work showing that responses on the attitude and desire subscales can be primed suggests they also measure state-like aspects of sociosexuality (Moss & Maner, 2016). The lack of evidence for effects of hormonal status on women’s desire for uncommitted sexual relationships is also arguably problematic for the theory that women are more likely to seek uncommitted sexual relationships with high-quality mates during the high-fertility phase of their menstrual cycle (see Gildersleeve et al., 2014 and Wood et al., 2014 for metaanalyses drawing opposite conclusions about how robust the evidence for this proposal is). Some versions of this theory predict an effect of hormonal status on women’s desire for uncommitted sexual relationships (Penton-Voak et al., 1999).

Arslan et al. (2017) recently reported that fertility had similar positive effects on partnered women’s reported in-pair and extra-pair sexual desire. This pattern of results is consistent with the effects of hormonal status on general sexual desire, but not sociosexual orientation, that we observed in the current study. We note here that other researchers have suggested that partnered women show hormone-linked changes in sociosexuality only if their partner is relatively unattractive (Gangestad et al., 2005). While it is unlikely that the partnered women in our study will predominantly have had partners who were more attractive than average, we do not rule out the possibility that considering partner or relationship characteristics could yet reveal hormone-linked changes in sociosexuality in some partnered women.

In conclusion, our analyses of a much larger dataset than those used in previous studies showed strong support for the proposal that changes in hormone levels, and progesterone in particular, are related to changes in women’s general sexual desire (Roney & Simmons, 2016). By contrast with other recent work (Grebe et al., 2016), however, we found no evidence that changes in women’s desire for uncommitted sexual relationships were related to changes in their hormonal status. Our results highlight the importance of employing large sample sizes to test hypothesized relationships between changes in women’s hormone levels and changes in their mating psychology.

## Acknowledgments

We thank Jim Roney, Ruben Arslan, Julia Junegner, and Aaron Lukaszewski for comments.

